# Benchmarking spatial interpolation methods for brain maps

**DOI:** 10.64898/2026.04.27.721143

**Authors:** Yigu Zhou, Vincent Bazinet, Bratislav Misic

## Abstract

The human brain is a unique biological space that hosts complex processes unfolding at multiple scales. To study these processes, an abundance of imaging technologies evolved over many decades to produce large-scale, dense mappings of structural and functional features. In parallel, a rich universe of techniques for cellular and molecular biology supplies us with fine-scale, highly specific and reliable measurements in sparse tissue samples. To represent cortical processes integratively across scales, spatial interpolation is necessary for bridging dense and sparse data. The absence of a field consensus for realistic interpolation of features over the whole brain prompts a comprehensive comparison of existing frameworks from the broader scientific literature. Here, we benchmark the performance of multiple deterministic (Inverse Distance Weighting, K-Nearest Neighbours, and Radial Basis Function) and stochastic (Spatially-Weighted Regression, Ordinary Kriging, and Regression Kriging) strategies first with simulated or empirical ground truths. We then demonstrate two use cases with *de novo* sparse brain data (intracranial EEG and microarray gene expression). In these experiments, we investigate how differences in data characteristics, such as spatial dependency structure and sampling distribution, impact the performance of different interpolation methods. Throughout the results, we consistently find that maps interpolated through spatially-informed stochastic frameworks such as Ordinary Kriging and Regression Kriging are more accurate and biologically realistic across geometric constraints, data modalities, and sampling conditions. This invites continued development of spatially-informed statistical frameworks for analyzing brain data and, more fundamentally, the biological processes that produce them.

## INTRODUCTION

The brain is an integrative system embedded in a meshwork of cells that communicate with each other, shaping our cognitive and behavioural repertoire. This biological cosmos is fundamentally built from a molecular blueprint specified by genetic programs, which are translated to proteins that define cellular units and wiring properties. Technological innovations in bio-imaging and recording enable access to a wide array of observations of brain structure and function with unprecedented precision. *In vivo* imaging techniques, such as magnetic resonance and nuclear imaging, generate dense whole-brain maps of anatomy (e.g., grey matter density, myelination) and dynamics (e.g., electromagnetic oscillations, hemodynamics, glucose metabolism).

In contrast, finer scale measurements use sparse *ex vivo* or invasive samples that are spatially restricted by availability of specimens or instrument access. To this day, data from brain tissue samples are the gold standard in neuroscience^26^, and there are increasingly refined methods to take measurements from tissue at the molecular and cellular resolutions. These expertly collected and curated datasets are now publicly available through large data repositories, promoting synthetic endeavours that integrate neurobiological insights across spatial scales^32,57^. For example, using *post mortem* tissue, spatially-resolved transcriptomics measure gene expression levels^35^; autoradiography measures neurotransmitter receptor density^102,103^; histology and cell staining map cellular density (popular empirical datasets include BigBrain^4^, NextBrain^13^, and BigMac^39^); antibody-based imaging maps synapse types and distributions^101^, as well as protein profiles (Human Protein Atlas)^85^; mass spectroscopy elucidates lipid profiles^68^; spectrophotometry tracks enzymatic activity and mitochondria phenotypes^59^. Via surgical access, intracranial electrophysiology generates direct recordings of parenchymal activity^25^. The integration of these fine-scale insights with large-scale features can help achieve a more comprehensive neurobiological cartography. To do so, spatial interpolation is the vital bridge between dense and sparse data.

The problem of spatial interpolation boils down to finding a function unifying a spatial process over discretely observed spatial locations that can be used to approximate values continuously across the whole spatial domain, including at unobserved locations. Although there exist infinite solutions to the problem, commonly encountered spatial interpolation models are invested with concepts from spatial statistics, as well as domain heuristics. Broadly speaking, spatial interpolation methods fall into two main categories: deterministic or stochastic^58,62^. The former category includes methods that assume homogeneous spatial variation within fixed neighbourhoods, and the latter category includes methods that either explicitly model patterns of variation across the spatial domain, or utilize adaptive neighbourhoods. Since those methods were initially developed for and routinely used with geospatial data^44,48,49,73^, the behaviour of different interpolation strategies within the confines of brain anatomy remains unknown. Spatial interpolation is a highly dynamic topic of research in disciplines that deal with spatial data such as geology and environmental sciences.

Brain data is spatial data, often represented as values mapped to a set of coordinates for which different standardized reference systems have been devised over decades of brain mapping research^51^. However, the application of spatial interpolation to brain data for densification of sparse maps both across and within data modalities currently lacks benchmarks and methodological consensus. One notable example of sparse brain data being interpolated differently is the Allen Human Brain Atlas (AHBA) microarray gene expression dataset^35^, which is a landmark dataset for imaging transcriptomics^24^. Since its publication, the sparsely sampled microarray data were interpolated differently to a dense mesh representation of the cerebral cortex using three-dimensional Ordinary Kriging^29^, K-nearest neighbours interpolation^53^, or K-nearest neighbours interpolation combined with kernel smoothing^94^. Among these *ad hoc* method choices, there is no unifying framework to assess accuracy and biological relevance of different approaches, as the spatial structure of brain data remains poorly understood. Moreover, a rich universe of highquality sparse brain annotations, beyond the AHBA, remains to be explored. Therefore, next-generation multiscale neuroscience requires a systematic investigation and comparison of spatial interpolation models by which to both address the practical need to compute dense maps, and the theoretical need to better understand the spatial structure of brain data.

Here, we select six published spatial interpolation methods that implement a deterministic (Inverse Distance Weighting, K-Nearest Neighbours, and Radial Basis Function) or stochastic (Spatially-Weighted Regression, Ordinary Kriging, and Regression Kriging) framework (Table 1, Fig. 1), and comprehensively compare their behaviour using simulated and empirical data. First, we use a harmonized ranking scheme to record relative performances using controlled simulations of spatial structure and sampling regime. Next, we use open datasets and apply each interpolation method to the same analysis with multiscale, multimodal empirical brain maps. Finally, we demonstrate two use cases where we interpolate *de novo* dense maps from sparsely sampled intracranial EEG (iEEG) and microarray data, and put them into biological context with densely sampled MEG and PET proxies. Although we focus mostly on neuroscience data and questions, to our knowledge this type of systematic benchmarking of spatial interpolation methods has not been performed in any other literature, so the results presented here are potentially informative to the broader natural sciences literature.

**Table 1.** Spatial interpolation methods. Overview of spatial interpolation methods for three-dimensional point sets considered in the reported analyses. The Description column indicates the primary point of methodological divergence for each framework. The Parameter column indicates pertinent model parameters to be either user-specified or estimated with sample data. Code and references to code for the implementation of all frameworks is available at https://github.com/netneurolab/zhou_interpolates.

| *Method* | *Description* | *Parameter(s)* |
| --- | --- | --- |
| **Deterministic** |  |  |
| *idw* Inverse Distance Weighting | Inverse distance-weighted average of values at known points to the π-th power | $\pi$ |
| *knn* K-Nearest Neighbours | Inverse distance-weighted average of values at $k$ nearest neighbours | $k$ |
| *rbf* Radial Basis Function | Kernel function-weighted average of values at $k$ nearest neighbours | $k$ |
| **Stochastic** |  |  |
| *swr* Spatially-Weighted Regression | Point-wise multiple linear regression from $m$-many covariates $X_m$ measured within bandwidth $b$ | $X_m, b$ |
| *ok* Ordinary Kriging | Gaussian process regression over the spatial domain from variogram $\gamma$ fitted at known points | $\gamma$ |
| *rk* Regression Kriging | Sum of spatially-agnostic multiple linear regression from $m$-many covariates $X_m$ measured at known points followed by Kriging with variogram $\gamma$ of regression residuals | $X_m, \gamma$ |

**Figure 1.**
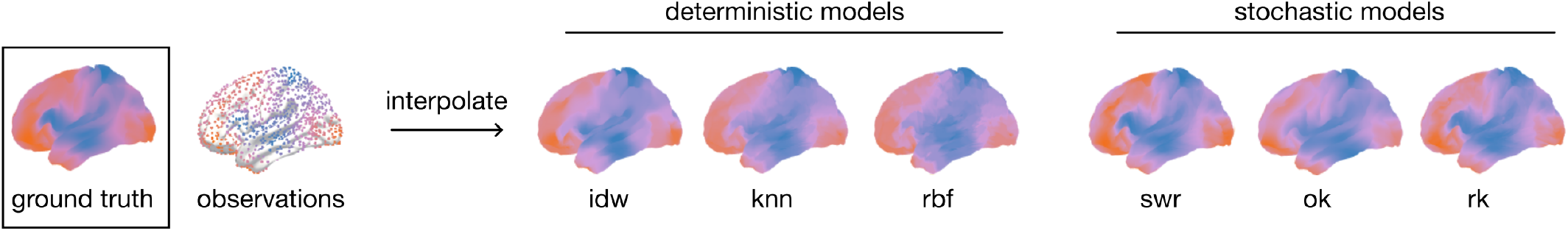
Spatial interpolation examples. It is impossible to observe a spatial process everywhere over a continuous spatial domain. Given a limited coverage of the spatial domain, spatial interpolation attempts to answer: what might be happening at the unobserved locations, based on the available observations? We show examples of the spatial interpolation applied to toy data on the brain. The spatial pattern that we specify as the “ground truth” is a realization of a Gaussian Random Field with the range parameter set to 45mm over the fsLR midthickness mesh. The “observations” are randomly taken from 50% the mesh. All deterministic interpolations are conducted with the same neighbourhood for comparison between their outputs. Regression models include the same five covariates transformed from the “ground truth” map. Kriging is done with a parametric Gaussian variogram fitted to the “observations”. See Methods subsection *Spatial interpolation frameworks* for more information.

## RESULTS

### Interpolating simulated spherical maps

We begin by examining the behaviour of different interpolation methods in the context of canonical spherical geometry using simulated data. Random spatially autocorrelated data are generated using gaussian random fields defined over a dense spherical mesh (fsLR, 4002 vertices)^64^. This approach involves the spectral randomization of a Gaussian variogram. The range of the variogram is set to 25-100mm to produce increasingly spatially autocorrelated maps. The maps are sparsely sampled at 5-50% of mesh vertices, either evenly or randomly (Fig. 2a, b). Toy data are constructed from simulated instances to serve as regression covariates for Spatially Weighted Regression and Regression Kriging interpolation (Fig. S1a). In this section, we explore the spatial structure of spatially autocorrelated maps and test how range and sampling density affect interpolation performance.

**Figure 2.**
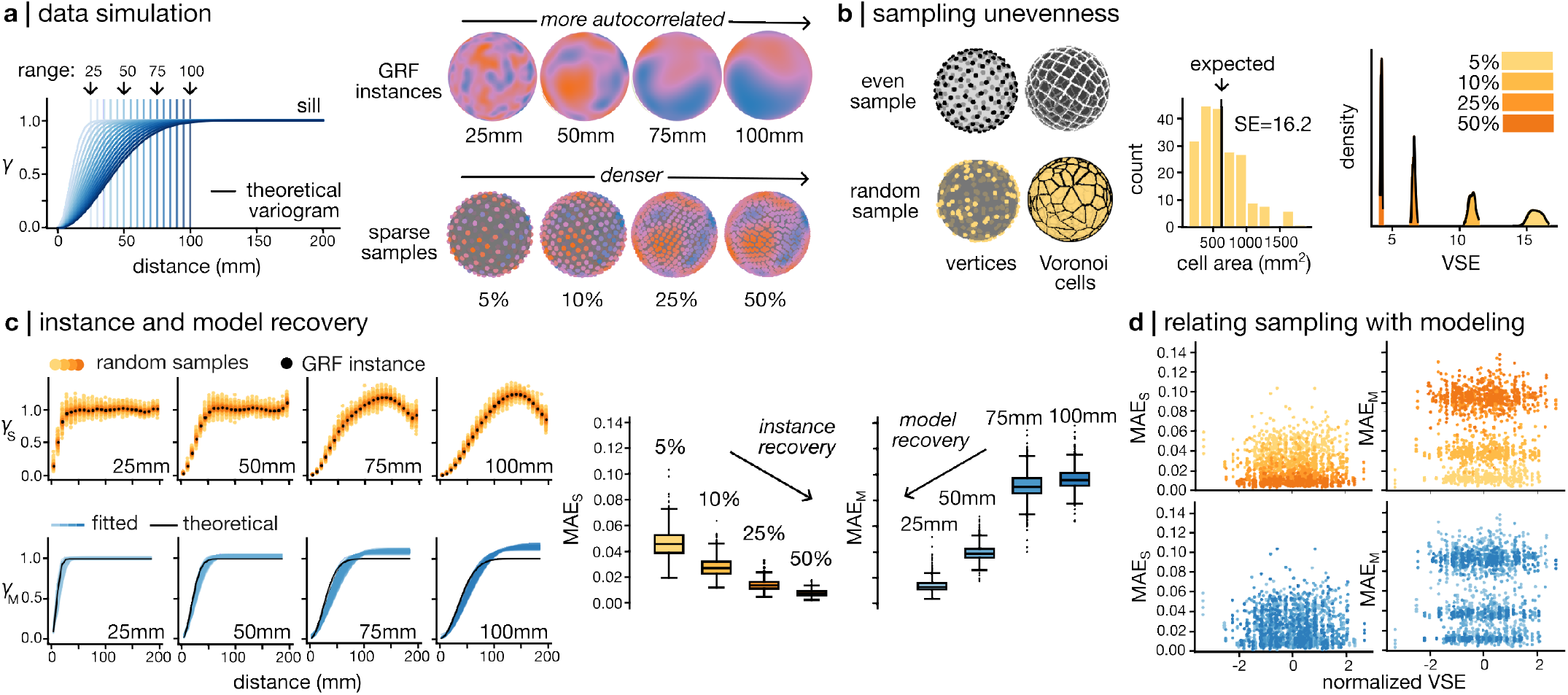
Mapping the spatial structure of simulated spherical data. **(a)** Spatial random fields are calculated from a theoretical Gaussian variogram model that has for parameters the sill, the range and the nugget. The sill of the variogram is the maximal distance-related semivariance (*γ*) value. The range of the variogram is the distance (in mm) at which *γ* plateaus. The nugget is the y-intercept, set to zero here. A series of dense maps with increasing spatial autocorrelation is computed from Gaussian variograms with unit-sill (variance of 1) and range set to 25-100mm. Sparse samples are taken from 5-50% of the mesh. Even samples are shown here to illustrate the extent of spatial coverage. **(b)** Unevenness is measured by comparing the distribution of vertex-wise surface areas of random samples to that a same-sized reference even sample. The standard error (SE) scores the variability of surface areas of Voronoi cells centered at random sample vertices with respect to the “expected” surface area from even Voronoi cells. We refer to this metric as the Voronoi standard error (VSE). The example shown here samples the mesh at 5%. From 100 repetitions of random sampling at different percentages, we examine the distribution of VSE across sampling schemes (*V SE*_50%_ = 4.17±0.05; *V SE*_25%_ = 6.63±0.10; *V SE*_10%_ = 11.0±0.24; *V SE*_5%_ = 15.8±0.48). **(c)** Experimental variograms from 100 random samples per range are compared against the full GRF experimental variogram (*γ*_*S*_). Instance recovery is measured by mean absolute error *MAE*_*S*_. Sampling at higher percentage produces a better recovery of the GRF instance variogram (lower MAE) across ranges. All sample experimental variograms are fitted to a parametric Gaussian model, and compared against the theoretical Gaussian variogram (*γ*_*M*_). Model recovery is measured by mean absolute error *MAE*_*M*_. Sampling from a shorter range produces a better recovery of the Gaussian variogram model (lower MAE) across sampling densities. **(d)** VSE normalized to sample size is related to instance (Adjusted *R*^2^ = 0.79, *F*_(*df*=13)_ = 3233, *p* = 0.00) or model recovery (Adjusted *R*^2^ = 0.94, *F*_(*df*=13)_ = 1643, *p* = 0.00) using a linear mixed effects model, with range and sampling percentage as random effects. Instance and model recovery both are significant and stably related to VSE within sample size or data range, but interaction effects between sample size and data range are not significant.

Sparse samples provide all the information for interpolation, so we are first interested in how variation in sampling produces different data. Since we are sampling randomly on the sphere, the absence of constraints makes it inevitable that random coverages would be uneven. We quantify unevenness by the surface area occupancy of vertices. For each randomly sampled set of vertices, we generate a Voronoi tessellation over the spherical mesh using the sample vertices as centroids^12^. The spatial coverage of each vertex is represented by the surface area of the Voronoi cell at that centroid vertex. A set of vertices that evenly samples the spherical mesh would have equal-sized Voronoi cells, or even surface areas, such that for a set of vertices of any size, there exists a unique value as expected Voronoi cell area. We represent the unevenness of spatial coverage as the standard error of Voronoi cell area distributions from the expected Voronoi cell area, henceforth referred to as “Voronoi standard error” (VSE). A larger VSE reflects a more uneven sampling scheme.

The absolute VSE of random samples increases with sampling sparsity (Fig. 2b). The spatial structure of sparse data is encapsulated by its variogram, which is a representation of the rate of change in dissimilarity over distance. Experimental variograms of random samples resemble the shape of theoretical Gaussian variograms at lower effective ranges by retaining the plateau of the partial sill of the variogram beyond the effective range. However, at higher effective ranges, random samples produce experimental variograms with a cyclical pattern beyond the effective range (Fig. 2c). Calculating variograms with geodesic distance so to accommodate for curved geometry yields similarly-shaped variograms (Fig. S2a). That being said, since parametric variogram models are derived in Euclidean space, without guaranteed compatibility with other definitions of distance^6^, we compute all distances in this report as Euclidean distances (Fig. S2b).

We then ask: how similar is the spatial structure of sparse samples to that of the ground truth dense data? And how well can sparse samples recover the underlying spatial process via modeling? To describe the recovery of the spatial structure of a GRF instance by sub-samples, we compute the deviation of experimental variograms taken from random samples (*γ*_*S*_) from the experimental variogram taken from the full GRF instance, separately at each range. To describe the recovery of the theoretical spatial structure of a GRF instance by modeling based on subsamples, we fit the experimental variograms to a Gaussian model (*γ*_*M*_) and compare fitted variograms to the theoretical variograms after sill-standardization. We use the Gaussian variogram model here because that is the model used to instantiate random fields. Instance and model recovery are optimal with a denser sample, at a shorter range, respectively (Fig. 2c). In other words, more accurate interpolation is expected from denser samples of a spherical GRF that has a more local profile of spatial autocorrelation. Instance and model recovery are anticorrelated (*R* = −0.83) to each other but not correlated to sampling unevenness (*R* = 0.013, 0.025; Fig 2c, d).

After exploring a diversity of sampling solutions and how they relate to the original data, we submit the various subsamples to interpolation over the entire spherical mesh, and assess the interpolatability of different conditions of range and sampling percentage. The absolute performance error of each method is scored as the root mean squared error (RMSE) between the interpolated map and the ground truth. We construct learning curves for GRFs of different ranges at increasing sampling density in percentage and analyze the behaviour of each model (Fig. 3a, Fig. S3b). GRFs with longer ranges are reconstructed with less error overall. Within the same range, reconstruction improves with a larger sampling percentage. The learning curve is steepest for GRFs of short ranges before plateau. Under the range of 35mm, the prediction error plateau does not reach below *RMSE* = 0.10 (99% variance explained). Among deterministic models, K-Neareast Neighbours interpolation is capable of approximating GRFs more accurately with less samples than Inverse Distance Weighting and Radial Basis Function. Among stochastic models, Ordinary Kriging produces similarly shaped learning curves but is by far the most optimal interpolator capable of reconstructing almost perfectly the original data with, in the best cases, less than 10% of sample vertices. Exceptionally, Spatially-Weighted Regression and Regression Kriging both perform relatively stably regardless of the condition of the data or sampling, with Regression Kriging (*RMSE* = 0.10-0.33) consistently outperforming Spatially-Weighted Regression (*RMSE* = 0.29-0.60). Overall, we find that individually, each spatial interpolation method is differently sensitive to the spatial autocor-relation structure of the data as well as sampling density.

**Figure 3.**
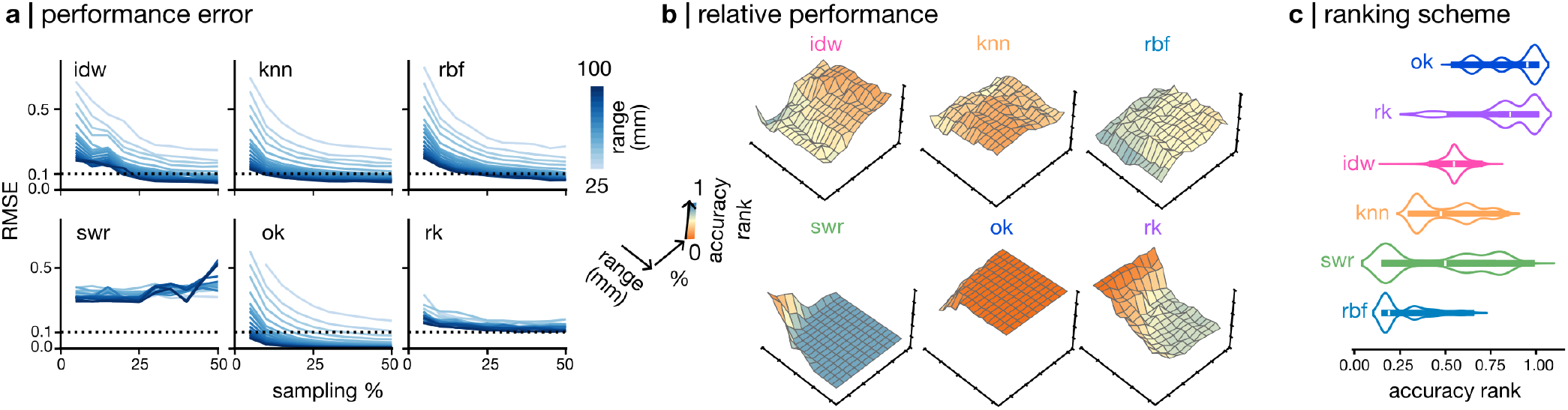
Interpolating simulated spherical maps. **(a)** Absolute performance of spatial interpolation is scored as the root mean squared error (RMSE). Here we show learning curves from even samples taken from 5-50% of the sphere. GRF instances of range 25-100cm are evaluated separately. The dotted line at *RMSE* = 0.1 marks 99% variance explained. **(b)** Relative performance of spatial interpolation aggregates across multiple metrics of similarity. Results from unique, even samples are shown here. Colour of each face on the surface maps represents the value of the accuracy rank score as well as the position on the vertical axis. **(c)** The ranking scheme pools together relative performance from random subsamples across ranges and sampling densities.

Next, we are interested in how each spatial interpolation method compare when submitted to the same task with the same data. We score the relative performance of different methods by ranking them across multiple metrics that assess similarity between the interpolated and the ground truth maps (Fig. 3b, Fig. S3b). The relative performance maps across range and sampling percentage suggest that short-range sparse samples (range under 40mm and sample size under 10% of the mesh) are more adequately modeled by Regression Kriging and Spatially-Weighted Regression, than by Ordinary Kriging and the deterministic models. In all other cases, Ordinary Kriging prevails as the optimal interpolator for GRFs.

The performance of spatial interpolation is known to be affected by data characteristics^49^. For the interpolation methods tested here, we expect data characteristics such as instance recovery and sampling unevenness to influence the performance of all methods similarly, and model recovery to influence the performance of kriging methods. To verify this, we use a linear mixed effects model to probe the relationship between accuracy rank and data characteristics, with method choice, range, and sampling percentage as random effects (Adjusted *R*^2^ = 0.606, *F*_(*df*=24)_ = 772.0, *p* = 0.00). Method choice is the dominant determinant of accuracy rank, with Ordinary Kriging ranking highest across data and sampling conditions, followed by Regression Kriging, Inverse Distance-Weighting, K-Nearest Neighbours, Spatially-Weighted Regression and Radial Basis Function (Fig. 3c). In addi-tion, instance recovery 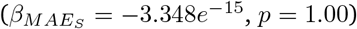 and sampling unevenness (*β*_*VSE*_ = −0.002, *p* = 0.665) do not significantly affect accuracy rank, supporting that input data characteristics affect all methods similarly. However, at a shorter range and with sparser samples, interpolation methods which include a regression step (Regression Kriging and Spatially-Weighted Regression) outperform Ordinary Kriging (Fig. 3b, c). Specifically for Regression Kriging, we hypothesize that in those conditions, since model recovery is worse, the additional regression step potentially provides an advantage over Ordinary Kriging at high variogram fit error provided regression covariates that are highly correlated to the independent variable (Fig. S1a). Using a linear mixed effects model, we relate accuracy rank for Ordinary Kriging and Regression Kriging to model recovery, with method choice, range, and sampling percentage as random effects (Adjusted *R*^2^ = −0.002, *F*_(*df*=15)_ = 0.5884, *p* = 0.886). Contrary to our hypothesis, we find that model recovery is not related to the overall high-ranked performance of Ordinary Kriging and Regression Kriging.

Taken together, we show that with GRF simulations, range and sampling density affect instance and model recovery without necessarily affecting relative spatial interpolation performance across ranges and sampling density. Indeed, method choice is the main determinant of relative performance by ranking. The ranking scheme consistently highlights methods that involve variogram modeling as more accurate for reconstruction of a GRF.

### Interpolating multiscale, multimodal brain maps

To investigate how interpolation methods behave in the context of brain anatomy, we use empirically collected group-averaged maps of neurobiological features^54^ that include gene expression (microarray profiling), structure (MRI), function (fMRI), neural dynamics (MEG), and metabolism (PET) (Fig. 4a). All of these feature maps are co-registered to the fsLR midthickness template excluding unmeasured vertices of the medial wall. To compare interpolation accuracy across methods, we take 100 subsamples of all feature maps at 5, 10, 25, and 50% using an anatomy-first approach where point sets are distance- and size-constrained by the Desikan-Killiany segmentation atlas^17^, mimicking the way human brain specimens are procedurally dissected along anatomical landmarks for measurement^95^ (Fig. 4b). Regression covariates for Spatially-Weighted Regression and Regression Kriging are selected as the top five feature maps most correlated to the target feature map (Fig. S1b). In this section, we explore the spatial structure of brain feature maps and test how spatial modeling and sampling density affect interpolation performance.

**Figure 4.**
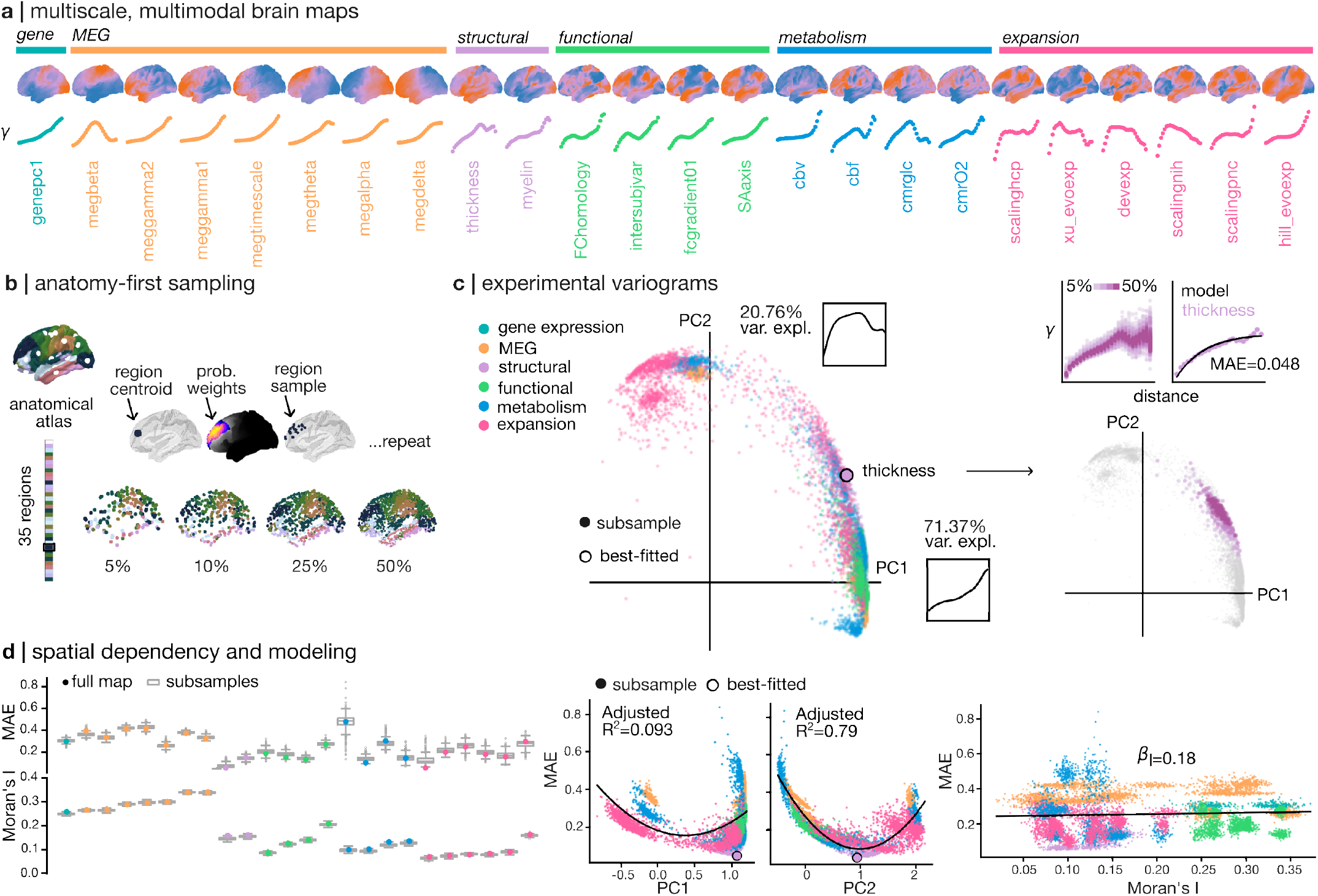
Spatial structure of brain maps. **(a)** Empirical dense maps that describe multiscale neurobiology are sourced from publicly available multimodal datasets, and co-registered to the canonical fsLR midthickness mesh. Experimental variograms of the original maps are computed over even distance bins. These empirical maps are sparsely sampled 100 times using an anatomy-first approach at 5-50% of the cortical mesh. **(b)** Anatomy-first sampling uses the Desikan-Killiany anatomical atlas which includes 35 contiguous brain regions of different sizes delineated by gyrification. Geodesic centroid of each region is defined as the center of a Gaussian kernel that covers the region. The kernel function assigns a probability weight to each vertex of the region. A region sample is proportional to the size of the region, and includes vertices chosen probabilistically based on distance from the region centroid. This is repeated for every region, and the region samples are assembled into a whole-brain sample. **(c)** Functional principal component analysis (FPCA) of experimental variograms of sparse samples highlight two main shapes for the experimental variogram via the first two principal components (PC1: 71.37% variance explained; PC2: 20.76% variance explained). Experimental variograms are fit to an exponential model and scored for fit error using mean absolute error (MAE) after sill-standardization. The best-fitted variogram is cortical thickness (*MAE* = 0.048). The spread of experimental variograms from cortical thickness subsamples are embedded in FPCA space in a spread-out fashion according to their shapes that vary based on sampling percentage. **(d)** Error of fit to an exponential model is different for each feature map. We relate MAE to PC1 (Adjusted *R*^2^ = 0.093) and PC2 (Adjusted *R*^2^ = 0.79) as positive quadratic functions. We assess the spatial autocorrelation of sparse samples using the global Moran’s I statistic. Spatial autocorrelation is also specific to feature maps. Spatial autocorrelation and exponential variogram model fit are related to each other with a mixed linear effects model, with range and sampling percentage as random effects (Adjusted *R*^2^ = 0.93, *F*_(*df*=53)_ = 7718, *p* = 0.00). Moran’s I and MAE are not overall related (*β*_*I*_ = 0.18, *p* = 1.27*e*^*−*8^ after Bonferroni correction), but are differently related within features, coloured by category.

The spatial structure of each feature map is summarized by an isotropic variogram across all sample vertices over 25 even distance bins (Fig. S2b). Each feature produces a differently shaped experimental variogram (Fig. 4a, Fig. S4a). We identify dominant variogram shapes by decomposing the experimental variograms from individual feature maps and their subsamples using functional principal component analysis (FPCA; Fig. 4c, Fig. S4a). The first component (PC1; 71.37% variance explained) captures a monotonic increase in semivariance as a function of distance. The second component (PC2; 20.76% variance explained) captures a rapid rise to a plateau at the maximal semivariance followed by a dip at the most distant bins. Subsamples of the same feature occupy an expanse of the FPCA space even though they form a cluster. This means that both the data itself and the sampling scheme affect the shape of the experimental variogram (Fig. 2c, Fig. S4a).

Given the variety of shapes of experimental variograms, we choose to describe them using a parametric exponential variogram model following existing literature reporting that spatial autocorrelation decays exponentially in the brain^34,47^. The best-fitted experimental variogram (*MAE* = 0.048) is computed from cortical thickness sampled at 25% (Fig. 4c, d), and the worst (*MAE* = 0.84) is computed from cortical blood volume sampled at 5%. The fit error (MAE) is related to the first and second components by quadratic functions (MAE-PC1: Adjusted *R*^2^ = 0.093, *RMSE* = 0.104, *y* = 0.19*x*^2^ − 0.14*x* + 0.18; MAE-PC2: Adjusted *R*^2^ = 0.79, *RMSE* = 0.050, *y* = 0.18*x*^2^ − 0.36*x* + 0.27) (Fig. 4d). Specifically, the second component provides an axis that centers the most exponential-like variogram, while higher error values spread out on either side of the PC2 median.

We next investigate whether the wide spread of model fit error reflects the inherent spatial dependency of the feature maps (Fig. 4d). For all feature maps included, spatial autocorrelation is measured using the Moran’s I statistic. The significance of spatial dependency is tested against 1000 permutation samples (Fig. S4b). We relate spatial autocorrelation to variogram fit error, with feature map and sampling parameters as random effects (Adjusted *R*^2^ = 0.93, *F*_(*d*=53)_ = 7718, *p* = 0.00). Across feature maps and sampling strategies, variogram fit error is overall weakly related to spatial autocorrelation of sampled data (*β*_*I*_ = 0.18, *p* = 1.27*e*^*−*8^ after Bonferroni correction; Fig. 4d). In other words, the brain-embedded feature maps clearly harbour a strong spatial dependency which is more complex than can be summarized by an exponential model.

How do spatial interpolation methods perform with the complex spatial pattern of brain feature maps? To assess the interpolatability of different feature maps, we subsample and interpolate each brain feature separately. The absolute performance error of each method is scored as the normalized root mean squared error (NRMSE) between the interpolated map and the original dense feature map. We construct learning curves for features of different categories. Across different features, the learning curve is steepest under 25% sampling density and shows limited improvement when the sample is doubled to 50%. For the majority of features, the prediction error can reach below *NRMSE* = 0.25 (94% variance explained) with between 25 to 50% of the mesh being sampled (Fig. 5a, Fig. S5a). However, only interpolation of the MEG maps produces less error than *NRMSE* = 0.10 (99% variance explained) (Fig. S5a). Overall, we find that it is possible to obtain an adequate reconstruction of a dense brain map with spatial interpolation with less than half of the original data.

**Figure 5.**
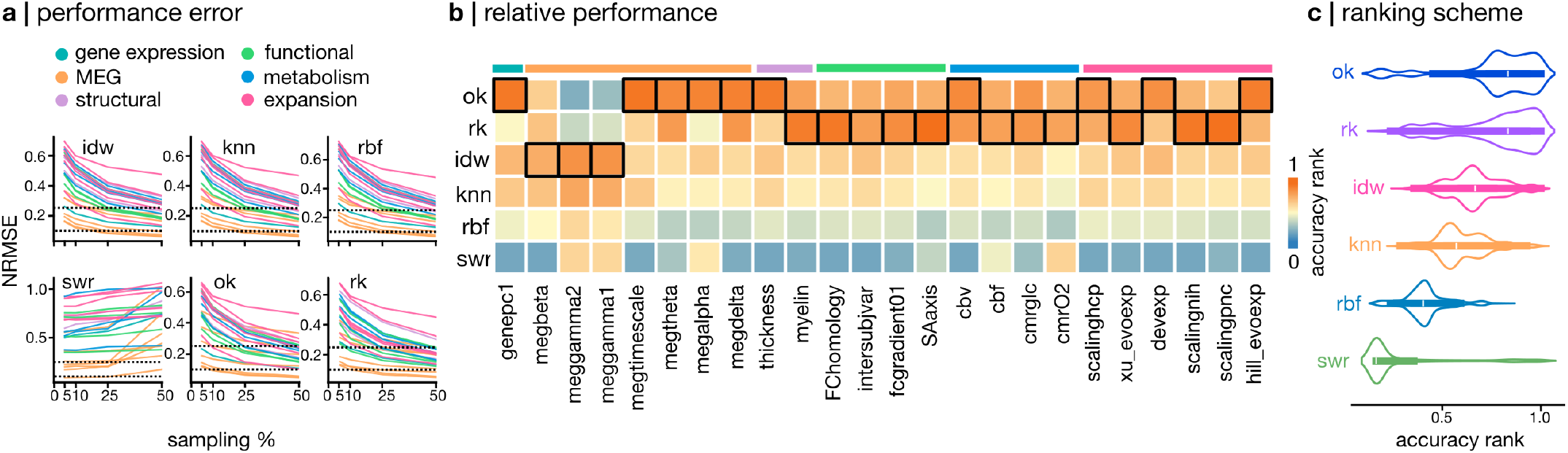
Interpolating multiscale, multimodal brain maps. **(a)** Absolute performance of spatial interpolation is scored as the normalized root mean squared error (NRMSE). Here we show learning curves from 100 anatomy-first samples taken from 5, 10, 25, and 50% of the brain surface, evaluated separately for representative features. The dotted lines at *NRMSE* = 0.25 and 0.1 mark 94% and 99% variance explained. Line colours correspond to feature category. **(b)** Relative performance via ranking is scored across empirical maps and averaged across sampling densities. The best-scoring interpolation method is boxed. **(c)** The ranking scheme pools together relative performance from all anatomy-first subsamples of all feature maps across sampling densities.

Since the success of interpolation varies based on the feature of interest, we create a feature-specific score board for each interpolation method. Relative performance of spatial interpolation methods are scored as an aggregate rank score from a selection of similarity metrics (Fig. S5a). The highest-ranked method across sampling regimes is most often Regression Kriging (for 11 features), followed by Ordinary Kriging (for 10 features) and Inverse Distance Weighting (for 3 features) (Fig. 5b). This tells us that despite variations in spatial modeling and interpolatability, kriging-related methods still prevail across vastly different data and sampling scenarios.

Nevertheless, to interpolate individual samples, we expect data characteristics such as spatial autocorrelation and modeling error to guide method choice. We use a linear mixed effects model to probe the relationship between accuracy rank and sample characteristics (variogram fit error via MAE and spatial autocorrelation via Moran’s I), with feature map and sampling parameters as random effects (Adjusted *R*^2^ = 0.56, *F*_(*df*=95)_ = 1363, *p* = 0.00). Similarly to our tests with simulated data, method choice is the dominant determinant of accuracy rank, with Ordinary Kriging ranking highest across feature maps and sampling conditions, followed by Regression Kriging, Inverse Distance-Weighting, K-Nearest Neighbours, Radial Basis Function and Spatially-Weighted Regression (Fig. 5c). Sample characteristics such as variogram model fit and spatial autocorrelation do not significantly impact the average ranking (*β*_*MAE*_ = 0.2, *p* = 0.19; *β*_*I*_ = −0.15, *p* = 0.119). We further investigate the circumstances in which Regression Kriging provides an advantage over Ordinary Kriging, by relating performance with variogram model fit error and spatial autocorrelation (Adjusted *R*^2^ = 0.57, *F*_(*df*=83)_ = 538.7, *p* = 0.00). At low variogram fit error, Regression Kriging outperforms Ordinary Kriging (*β*_*rk*_ = 0.20, *p* = 9.45*e*^*−*40^ after Bonferroni correction). At high variogram fit error, the performance of Ordinary Kriging decreases (*β*_*MAE*:*rk*_ = −0.49, *p* = 0.0 after Bonferroni correction). Different from spherical simulations, the regression step from Regression Kriging provides an advantage over Ordinary Kriging only when variogram model fit error is minimal. This difference could be accounted for by the diversity of variogram shapes in the empirical data and the use of empirically chosen regression covariates as opposed to synthetic ones. Nevertheless, provided adequate variogram modeling (*MAE <* 0.2), Ordinary Kriging and Regression Kriging outperform Spatially-Weighted Regression and deterministic methods.

Taken together, we show that empirical brain features are spatially-dependent and display a large variety of spatial structure. Sampling density of these dense feature maps change the shape of the variogram and affects the accuracy of an exponential variogram model. Nevertheless, kriging methods are most often the best-ranked methods across features. The ranking scheme also consistently highlights methods that involve variogram modeling as more accurate for reconstruction of a dense brain map. As such, the behaviour of spatial interpolation methods translates from spherical to brain geometry.

### Interpolating sparse brain data

In this section, we use the six interpolation frameworks introduced here on two openly available datasets of sparsely sampled brain data, and compute *de novo* dense maps. It is important to note that the upcoming experiment involves the interpolation of fine-scale neurobiological information to a dense brain mesh. This information is not available to dense sampling and therefore is fundamentally different from the large-scale features we examined in the previous sections. Due to the absence of a dense ground truth, we determine that *de novo* densified maps should account for both within-modality specificity and biological interpretability. Meeting these needs, we trial a benchmarking pipeline that separately assesses densified maps by criteria (i) correspondence to the original sparse data and (ii) correspondence to a dense proxy map obtained from a different modality. Specifically, we show use cases comparing alpha band (8-12Hz) power measured via intracranial electroencephalography (iEEG) or magnetoencephalography (MEG) and 5-hydroxytryptamine receptor 1A (HTR1A) density measured via microarray transcriptomics or protein binding from positron emission tomography (PET).

To produce dense maps of fine-scale neural dynamics, we use awake iEEG data from the Open MNI iEEG atlas which include timeseries from 1772 channels collected from 106 patients with focal epilepsy^25^. Channels that compose this normative iEEG atlas are selected based on strict criteria for non-pathological activity. We obtain the relative power of the alpha frequency band using the spectrogram of the timeseries and map the values to the position of each channel in MNI space, then interpolate them to the fsLR midthickness mesh. To validate the within-modality specificity of interpolation methods, we conduct five-fold cross-validation using an anatomy-informed and distance-dependent approach with the original channel coordinates (Fig. S6a).

We calculate experimental variograms for the full sparse iEEG data, the dense MEG data, and crossvalidation samples within even distance bins over the entire left-hemispheric iEEG coverage. The full sparse iEEG data vary according to distance in a roughly linear fashion (Fig. 6a, Fig. S6b). The experimental variograms of cross-validation samples are more similar to the full iEEG sample at shorter distance bins (Fig. S6b). Next, we fit an exponential variogram model to the full sparse iEEG data and gauge modeling error via MAE. Values for modeling error from cross-validation samples and from the full iEEG sample are within similar intervals (*p* = 0.49). To compare spatial covariance structure between iEEG and MEG data, we also fit an exponential variogram model to the full dense MEG data, binned according to iEEG coverage. The fitted exponential variogram in both cases show similar effective range estimates (*r*_*MEG*_ = 153.; *r*_*iEEG*_ = 150.), suggesting that the spatial autocorrelation radii for alpha bandpower are comparable across modalities. However, MEG-sampled alpha bandpower has 2.50-times higher overall spatial variance, which is consistent with large global changes due to smoothness of the MEG map (Fig. 6a). Overall, the spatial model for the same quantity (alpha band power) across biological scales is similar in patterns of local variations but disparate in the global smoothness. The disparity is expected and attributable to the difference in spatial resolution between iEEG and MEG.

**Figure 6.**
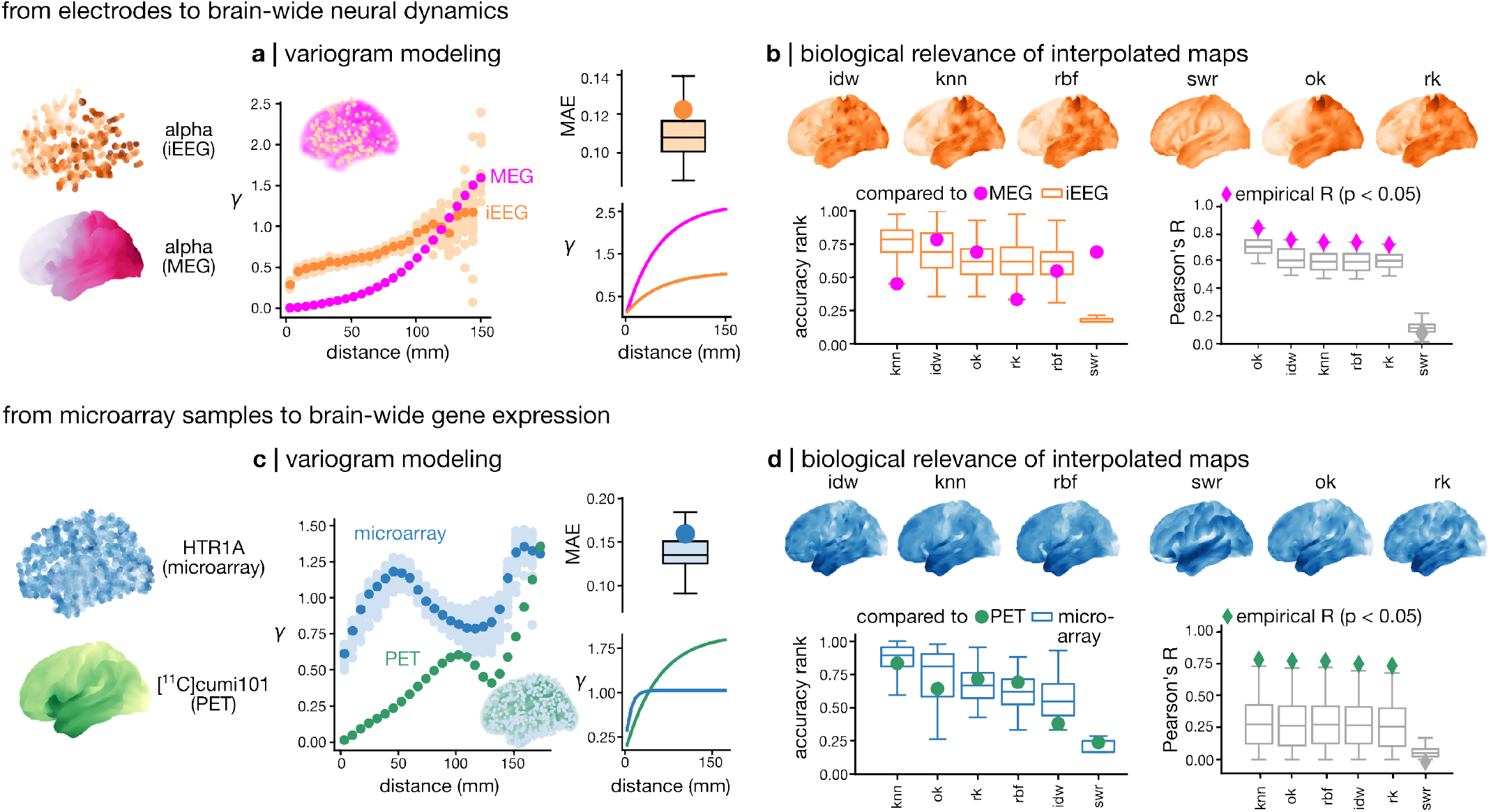
Interpolating sparse brain data. **(a)** We map alpha bandpower (8-12Hz) from sparsely sampled intracranial electroencephalography (iEEG) and from densely sampled magnetoencephalography (MEG). Experimental variograms are sill-standardized. The experimental variogram for MEG is scaled according to the MEG/iEEG sill ratio for visualization. We assess the adequacy of experimental variogram model using an anatomically informed distance dependent cross-validation approach. Mean absolute error (MAE) is calculated between the exponential variogram fitted over the full iEEG sample and experimental variograms from cross-validation iEEG samples (light orange) or the full iEEG sample (orange). We assess the cross-modal similarity of the spatial variation in alpha bandpower by comparing fitted parameters (effective range *r* and sill estimate *γ*) over the full MEG and iEEG samples (*r*_*MEG*_ = 153.; *r*_*iEEG*_ = 150.; *γ*_*MEG*_*/γ*_*iEEG*_ = 2.50). **(b)** We showcase the interpolated iEEG maps ranked by similarity to the MEG map. Accuracy ranks are scored for cross-validation iEEG samples against the full iEEG data (boxes) and for the full iEEG sample against the full MEG data (magenta). We quantify the similarity of interpolated iEEG maps with the MEG map via Pearson’s R while accounting for spatial autocorrelation. **(c)** We map 5-hydroxytryptamine receptor 1A (HTR1A) gene expression from sparsely sampled microarray transcriptomics data, and densely sampled [^11^C]cumi101 tracer binding from positron emission tomography (PET). Experimental variograms are sill-standardized. The experimental variogram for PET is scaled according to the PET/microarray sill ratio for visualization. We assess the adequacy of experimental variogram model using a anatomically informed distance dependent cross-validation approach. MAE is calculated between the exponential variogram fitted over the full microarray sample and experimental variograms from cross-validation microarray samples (light blue) or the full microarray sample (blue). We assess the cross-modal similarity of the spatial variation in HTR1A by comparing fitted parameters over the full PET and microarray samples (*r*_*PET*_ = 173.; *r*_*microarray*_ = 27.; *γ*_*PET*_ */γ*_*microarray*_ = 1.92). **(d)** We showcase the interpolated gene expression maps ranked by similarity to the PET map. Accuracy ranks are scored for cross-validation microarray samples against the full microarray data (boxes) and for the full microarray sample against the full PET data (green). We quantify the similarity of interpolated gene expression maps with the PET map via Pearson’s R while accounting for spatial autocorrelation.

By criterion (i), the interpolation method that best retains the patterns of the original iEEG data is K-Nearest Neighbours interpolation, followed by Inverse Distance Weighting, Ordinary Kriging, Regression Kriging, Radial Basis Function, and Spatially-Weighted Regression. By criterion (ii), the interpolation method that produces the best correspondence to the MEG reference map is Ordinary Kriging, followed by Inverse Distance Weighting, K-Nearest Neighbours, Radial Basis Function and Regression Kriging, all significantly so after accounting for spatial autocorrelation using null models (1000 iterations) generated using the Moran Spectral Randomization approach^93^ (Fig. 6b, Fig. S6c, d). Since Ordinary Kriging is favoured by both criteria, it is the most suitable method for interpolating iEEG data among those surveyed here.

To produce dense maps of fine-scale measurements of the HTR1A receptor, we use microarray bulk transcriptomics data from the Allen Human Brain Atlas and filter for probes with high differential stability (*>* 0.1), retaining values from 3468 tissue samples collected from six brains and mapped to the MNI space^35,53,100^. We note that microarray transcriptomics quantifies mRNA copies, while PET quantifies tracer binding to a protein. Therefore, correspondence is not guaranteed between the two measurements. However, HTR1A has been established to have adequate gene-to-receptor correspondence in our data and elsewhere^29,31,50^. We closely examine different interpolation methods for whole-brain expression of HTR1A receptor gene and compare it with the HTR1A receptor protein binding measured by PET tracer [^11^C]cumi101^9^. Cross-validation is applied to establish within-modality specificity same as above.

We calculate experimental variograms for the full sparse microarray data, the dense PET data and crossvalidation samples within even distance bins over the microarray sample coverage. The full sparse microarray data vary according to distance in a cyclical fashion (Fig. 6c, Fig. S6b). Next, we fit an exponential variogram model to the full sparse microarray data and gauge modeling error via MAE. Values for modeling error from cross-validation samples and from the full microarray sample are within similar intervals (*p* = 0.27). To compare spatial covariance structure between microarray and PET data, we also fit an exponential variogram model to the full dense PET data, binned according to microarray coverage. The fitted exponential variogram in both cases differ in effective range (*r*_*PET*_ = 173.; *r*_*microarray*_ = 27.) as well as sill estimates. PET-sampled receptor protein binding has 1.92-times higher overall spatial variance, which is consistent with large global changes due to the smoothness of the PET map. This means that the spatial models of HTR1A gene and protein expression are disparate. However, considering that with HTR1A data, variogram model fit error is on average higher than with alpha band data, the exponential model may not be well-suited for the complex pattern of HTR1A in the brain.

By criterion (i), the interpolation method that best retains the patterns of the original microarray data is K-Nearest Neighbours interpolation, followed by Ordinary Kriging, Regression Kriging, Radial Basis Function, Inverse Distance Weighting, and Spatially-Weighted Regression. By criterion (ii), the interpolation method that produces the best correspondence to the PET reference map is K-Nearest Neighbours interpolation, followed by Ordinary Kriging, Radial Basis Function, Inverse-Distance Weighting and Regression Kriging, significantly so after accounting for spatial autocorrelation using null models (Fig. 6d, Fig. S6c, d). The best correlation value is similar to previously published values (*R*_*knn*_ = 0.78)^31^. Since K-Nearest Neighbours is favoured by both criteria, it is the most suitable method for interpolating microarray gene expression data among those surveyed here.

## DISCUSSION

In the present work, we systematically examine the behaviour of six distinct and representative spatial interpolation methods from elemental categories, in the specific contexts of spherical GRF simulations, empirical brain surface maps and two use cases with empirical sparse brain maps. We also analyze the impact of spatial sampling strategy and data characteristics on the relative performance of each method by testing with a range of simulated and empirical data that express different degrees of spatial autocorrelation, and sparse samples of different sizes and sampling approaches (random, even, anatomy-first). Across all experimental contexts, the most successful dense map computation method evaluated by an aggregate accuracy score is Ordinary Kriging or Regression Kriging. Aside from specific scenarios where the parametric estimation of the spatial covariance structure captures the data poorly, stochastic methods generally outperform deterministic frameworks. Nonetheless, deterministic frameworks such as K-Nearest Neighbours and Inverse Distance-Weighting are safer choices when spatial modeling is suboptimal.

Throughout our results, we assess the relative performance as opposed to absolute performance. Spatial interpolation methods, when assessed across the same criteria, produce the consistent average ranking scheme summarized above. Indeed, it is expected that task difficulty penalizes all interpolation frameworks similarly, as they fundamentally differ in the set of assumptions each makes about input data that is baked into their design. Deterministic kernel-based methods reliably apply homogeneous rules at a consistent spatial scale to produce smooth maps that lack local variability and details. Hence, they reveal a consistent absolute performance across conditions that is often only slightly surpassed by kriging-related methods. Meanwhile, stochastic methods that apply parametric kriging models assume stationarity and isotropy that optimally reconstruct spherical GRFs. However, the complex geometry of the brain eludes these parametric models. In addition, the quality of spatial regression, either within Regression Kriging or in the style of Spatially-Weighted Regression, most importantly relates to the quality of regression covariates. In other words, a regression model evades the constraints of variogram modeling. When calibrated with appropriate covariates, Spatially-Weighted Regression can be more optimal than Kriging methods.

We also briefly examine those behaviours for spatial *extra*polation. That is, a special case of spatial prediction for samples *outside* the domain of observations. Cases of spatial extrapolation are simulated with anatomy-informed distance-dependent cross-validation samples, and produce relatively stable ranking of the different spatial interpolation methods examined here. The consistency of the ranking scheme across data types, sampling conditions, in both interpolating and extrapolating tasks suggests the existence of a potentially unifying rule of spatial patterning across the whole brain surface.

A key takeaway from the study of spatial structure and interpolation methods in our work is the necessity of an optimal spatial model both for the practical goal of computing dense maps, and the theoretical pursuit of modeling brain processes. According to Matheron’s theory of regionalized variables^56^, all spatial processes can be described by a combination of a deterministic and a stochastic component, describing an overlapping pattern of a global trend with local changes. This theory unifies kriging models and opened up for a virtually endless stream of optimized ways to compute spatial covariance functions. Although we limit ourselves, as a first exploration of spatial interpolation of brain maps, to parametric models^11,29,47^, more complex Bayesian approaches to spatial modeling that involve stochastic partial differential equations (SPDE) have been used in neuroscience to account for flexible and non-stationary spatial patterns^46,99^. However, issues of non-stationarity and anisotropy have not yet been considered in kriging interpolation for brain data. A natural extension of the present work is the investigation of more complex variogram models for more biologically realistic spatial simulations.

Moreover, we note that a common procedure of data postprocessing for multiscale neuroscience involves region-wise mappings according to a parcellation atlas, such as the Desikan-Killiany atlas used here. Parcellation, when applied to sparsely sampled data in the brain volume, involves averaging a potentially heterogeneous point set into a single value represented on unevenly sized polygons over the brain surface. This practice extensively coarsens highly spatially specific measurement, especially those observed at finer scales via *ex vivo* or invasive sparse sampling. In addition, potential biases introduced by averaging values in differently sized parcels have been noted as limitations in publications (e.g.,^53,94^). A possible avenue for addressing areal representation of inhomogeneous spatial information is areal interpolation, which calculates feature values for polygons by combining shape information with distance relationships^14^. We consider an exploration of areal interpolation strategies an interesting next step toward an enriched understanding of brain geometry.

In summary, this study presents a necessary independent comparison of representative spatial interpolation methods. We contextualize different methods in the same controlled settings to obtain a reproducible ranking scheme. Our results introduce benchmarking approaches that are intended as a resource for neuroscientists who work with sparse data. Simultaneously, the results are also a guide for those who are acquiring sparse data, as a starting point for devising optimal/economical sampling strategies. More broadly, our results invite further research in spatial models that can realistically account for both the general principles and local heterogeneity of brain cartography. Altogether, with the growth of open-data initiatives around the globe, our work provides foundational methodology for data integration while illustrating the relevance of spatial modeling for multiscale neuroscience.

## METHODS

### Spatial interpolation frameworks

In this section, we briefly describe spatial interpolation methods as they were initially proposed in the literature, then explain specific details of implementations used in the current report (Table 1). We also explain how parameter values are chosen for each model that we include in our comparative pipeline. Code used to run interpolation frameworks are adapted from publicly available interdisciplinary Python libraries for spatial statistics and freely available at https://github.com/netneurolab/zhou_interpolates.

The primary goal of spatial interpolation is to render continuous spatial processes from sparsely sampled data within a defined spatial domain. A point estimate can be expressed as

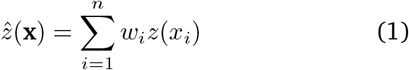

where the estimated value 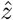 at location **x** is written as a weighted average across scalar values of a process *z*, observed across finite *n*-many sampled locations *x*_*i*_ (coordinate vectors). Different spatial interpolation methods diverge in the definition of the weights *w*_*i*_, which could be based on a deterministic kernel approximation, or a stochastic method that include spatially-conditioned coefficients or covariance function. We survey interdisciplinary literature for spatial interpolation methods for three-dimensional point data, which is the data format most commonly encountered in neuroimaging with both volumetric and surface-based mappings. We implement and test six exact interpolators (i.e., retaining values of sampled points) that follow deterministic (Inverse Distance Weighting, K-Nearest Neighbours, Radial Basis Function) or stochastic (Spatially Weighted Regression, Ordinary Kriging, Regression Kriging) frameworks.

#### Deterministic models

*Inverse Distance Weighting*. As the name implies, Inverse Distance Weighting (IDW) interpolation assumes a homogeneous distance relationship and returns spatial weights inversely proportional to distances to a power constant^81^. This implies modifying Equation 1 to

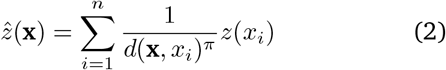

with power *π* and distance vector *d* between query point and sample points in Euclidean space. Our comparative pipeline chooses the optimal power constant by iteratively testing IDW models with *π* set to an integer between 2 and 20, and retaining the best ranking model. Neuroscientific applications of IDW include densification of cortical surface maps for visualization^16,30^, tissue classification^43^, or correction of spatially misaligned values^91^.

#### K-Nearest Neighbours

The K-Nearest Neighbours interpolation (KNN) algorithm relies on user-defined neighbourhoods^15,23^, and computes some point estimate as

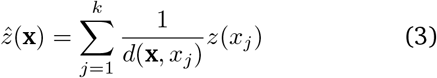

the inverse distance weighted average of *z* at some *k* ∈{1…*n* − 1} neighbours of **x**. Our comparative pipeline chooses the optimal number of nearest neighbours by iteratively testing KNN models with *k* set to an integer between 2 and 20, and retaining the best ranking model. This approach is implemented in the imaging transcriptomic toolbox abagen to compute transcriptomic gradients across the human cortex^53,94^.

#### Radial basis functions

The Radial Basis Function approach applies kernel function Φ to compute 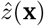

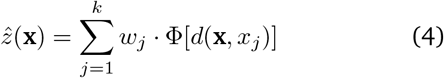

with *k* nearest neighbours from the sample. Weights are obtained by solving the linear system below.

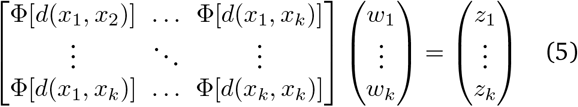

In this report, we select the Gaussian kernel function

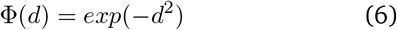

due to its popular use in the context of Random Field Theory applied to human brain mapping^97^. The interpolation function is used out-of-the-box from the `scipy` toolbox’s `RBFInterpolator` class.

#### Stochastic models

##### Spatially-Weighted Regression

What we refer to as Spatially-Weighted Regression (SWR) in this work is originally published under the name of “Geographically-Weighted Regression” in^10^. This is a multivariate regression approach that accounts for spatial relationships inside neighbourhoods. We test a version of SWR from^70,96^, implemented in the `mgwr` package^69^. Briefly, SWR builds on a traditional linear regression model

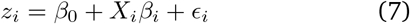

where *z*_*i*_ is a scalar attribute measured at point *i* ∈{1…*n*} which can be represented as a linear combination of a coefficients vector *β*_*i*_ with coordinates *x*_*i*_. The error term *ϵ*_*i*_ is normally distributed. The regression coefficients are spatially-weighted by estimating an *m*-elements vector

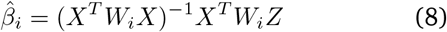

at point *i*. Here, *X* is a *n*-by-(*m* + 1) design matrix stacking together *m* regression covariates across sampled points and a leading column of ones for the intercept; *Z* is the vector of attributes across *n* sample points; *W*_*i*_ is a diagonal weights matrix specifying pairwise relationships between sample points *i, j* ∈ {1…*n*}. Our analyses use an adaptive bi-square kernel for spatial weights between points *i* and *j* based on their Euclidean distance *d*_*ij*_ such that

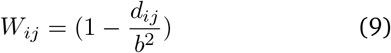

within a bandwidth *b* selected to minimize estimation error via corrected Akaike Information Criterion through a golden section search routine.

##### Ordinary Kriging

Kriging interpolation calculates spatial weights based on a model variogram relating pairwise distances to changes in attribute values. Ordinary Kriging (OK), the most widely used variation of Kriging, was used for neuroimaging data science in^29^. Spatial weights are related to the covariance function *C* in a system of equations

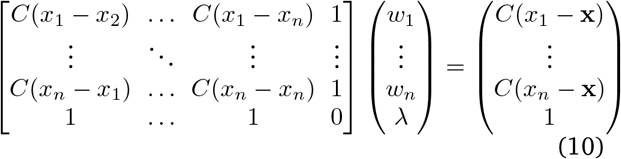

where *λ* is the Lagrange multiplier. The spatial covariance function *C* is related to variogram *γ* as

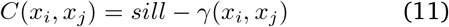

where the sill is defined as the maximum of *γ*. Assuming an isotropic variogram *γ* and stationarity of the mean, spatial weights are obtained by solving for *w*_*i*=1…*n*_, then applied to interpolation according to Equation 1.

#### Regression Kriging

The Regression Kriging method is a two-step routine which combines linear regression with univariate kriging^2,67^. We use the implementation of Regression Kriging from the pykrige package that first fits a Support Vector Regression model to scalar data, then performs Ordinary Kriging on the residuals with a variogram model *γ* fitted with the sampled data^61^.

### Performance evaluation

Sparse samples from simulated or empirical data in surface or volume space are interpolated to the entire spatial domain of a target surface (fsLR, 4002 vertices) based on Euclidean distances between vertex positions^89^. We compare interpolated and actual values at unsampled vertices, using an aggregate rank score *a* ranging from 0 to 1 that balances multiple metrics of accuracy (larger Pearson correlation coefficient, Spearman correlation coefficient, structural similarity index, and *R*^2^ values raise rank; larger root mean square error, Jensen-Shannon dissimilarity, and Kolmogorov-Smirnov distance values lower rank), as 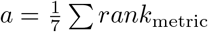. Reliability of perfor-mance for a random, or anatomy-first sampling regime is assessed by iteratively testing 100 different samples and taking the mean rank score across iterations.

### Variography

The experimental variogram captures the rate of change in some feature measure across distance. Pairwise distances from points on a surface mesh or a volumetric grid are collapsed into 25 even bins. Semivariance of feature values is calculated between pairs of points as

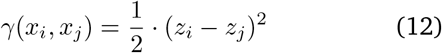

then averaged within each distance bin. This experimental variogram is fitted to either a parametric Gaussian model defined as

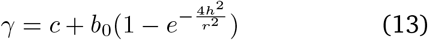

or exponential model defined as

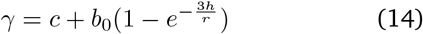

in Euclidean space, with *c* being the partial sill and the *b*_0_ being the y-intercept, or nugget, *h* being distance and *r* being range. We fit experimental variograms from simulated data on the spherical mesh to a Gaussian variogram model since that is the initiating model for data simulations. For empirical cortical data, we fit their experimental variograms to an exponential variogram model following existing descriptions of spatial dependencies in the brain^34,47^. We convert fitted variogram models for Ordinary Kriging and Regression Kriging using the pipeline implemented in `Scikit-GStat`^63^.

### Sampling unevenness

To characterize the unevenness of sparsely sampled point sets over a surface mesh, we compare the Spherical Voronoi diagrams of random or anatomy-first samples to those of a same-sized approximately even sample. First, we place a same-sized point set along a Fibonacci lattice on an ideal sphere radially equivalent and aligned to the discretized fsLR spherical mesh. This involves mapping the golden angle 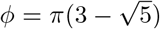 onto a sphere with unit radius with *i* ∈ {1…*n*} points represented by coordinates *α*_1…3_ along the x-y- and z-axes as

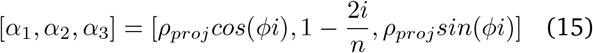

where the projection radius 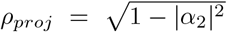 on a unit sphere. We then scale the unit coordinate vectors by radius *ρ* = 200mm. This approach offers an approximately equidistant sampling. We compute spherical Voronoi diagrams of the Fibonacci sphere sample and measure their mean surface area using the implementation for `SphericalVoronoi` from the Python `scipy.spatial` module^12^. This indicates the surface area coverage of sample points that is expected if the mesh were evenly sampled. The width of deviations from this expected value (standard error) is therefore an indicator of unevenness of sampling.

### Simulated spherical data

To produce spatially autocorrelated continuous data, we instantiate scalar maps over the fsLR canonical sphere using Gaussian random fields. We rely on the randomization approach implemented in GSTools^64^. Random maps are generated using Gaussian variograms with with sill of 1, nugget of 0, and range set to 25-100mm. This data simulation approach has been investigated previously to generate null models that account for spatial autocorrelation patterns in neuroimaging data science publications^7,11,55^.

Next, we sparsely sample 5-50% of the simulated maps using randomly generated point sets, or point sets evenly distributed over the mesh as approximated by a Fibonacci lattice. For random samples, GRF instances with ranges 25, 50, 75 and 100mm are each sparsely sampled at 5, 10, 25 and 50% with 100 different random point sets, creating a total of 4 × 4 × 100 unique interpolation cases per interpolation function. For even samples, GRF instances with ranges 25-100mm spaced by increments of five are each sparsely sampled at 5-50% spaced by increments of five, each with a unique Fibonacci sphere, creating a total of 16 × 10 unique interpolation cases per interpolation function.

### Empirical surface data

To examine the relevance of different methods for interpolating empirical data to cortical geometry, we selectively test them with feature maps of multiscale neurobiology from datasets in `neuromaps`^54^. We include 24 maps that are available in surface coordinate systems from six different biological categories including gene expression (gene PC1)^35^, MEG (alpha, beta, delta, low gamma, high gamma, theta power distribution, and intrinsic timescale)^79,80,90^, structure (myelin and cortical thickness)^28^, function (homology, gradient, inter-subject variability, and sensory-association axis)^52,60,84,98^, metabolism (cerebral blood volume, cerebral blood flow, metabolic rate of oxygen, metabolic rate of glucose consumption)^88^, and expansion (evolutionary cortical expansion, cortical areal scaling during development, and for the HCP, NIH, and PNC datasets)^37,72,98^. Each brain map is resampled to the fsLR midthickness mesh with multimodal surface matching^74,75^. These empirical maps display different global spatial autocorrelation as assessed by Moran’s I statistic

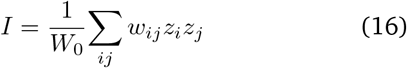

where *W*_0_ is the sum of all pairwise inverse Euclidean distances *w*_*ij*_ from a row-normalized distance matrix, and *z*_*i*_, *z*_*j*_ ∼ *N* (1, 0) are standardized scalar values from the feature maps.

### Empirical volumetric data

A repository of volumetric data including 30 positron emission tomography (PET) maps, 1 Single-photon emission computed tomography (SPECT) map and 1 arterial spin labeling (ASL) map is sourced from publicly available datasets available in the MNI152 template space at different resolutions. We include PET maps for the synaptic vesicle glycoprotein 2A^22^, metabolic glutamate receptor-5^19,33,82^, ionotropic GABA receptor A^20^, *α*2*β*4 acetylcholine receptor^38^, M1 acetylcholine receptor^65^, vesicular acetylcholine transporter^1,8,33^, serotonin receptor types 1A^78^, 1B^27,78^, 2A^78^, 6^71^, serotonin transporter^21,78^, norepinephrine transporter^18,36^, dopamine receptor type 1^41^, dopamine receptor type 2^3,40,83^, dopamine transporter^76^, opoid receptor type kappa^92^, opioid receptor type mu^42,86^, cannabinoid receptor type 1^45,66^, and histamine receptor H3^27^. We also include a SPECT map for the dopamine transporter^20^, and an ASL map that characterizes mean cerebral blood flow^77^. The maps are fetched through the neuromaps tool in the original template space in which they are available^54^. Then we compute a nonlinear transform for each map separately, mapping values from its original template space to the MNI ICBM152 nonlinear symmetric 2009c template using a quick nonlinear registration pipeline in Advanced Normalization Tools (ANTs) ecosystem^87^.

### Anatomy-first sampling of the cortical mesh

Empirical data are sparsely sampled at 5, 10, 25 and 50% of the brain surface using the Desikan-Killiany segmentation atlas as an anatomical prior^17^. We compute the geodesic centroid coordinates for each of the 35 contiguous regions of the atlas (medial wall included). A Gaussian kernel with sigma set to 0.5 is centered at the geodesic centroid of each atlas region, and used to probabilistically sample the region. The representation of vertices from each region is proportional to its size. We create a reduced midthickness mesh by removing medial wall vertices from the fsLR midthickness cortical mesh for the left cerebral hemisphere, retaining 3599 cortical vertices. This is used as sampling and interpolation mesh for empirical data analyses and use cases with iEEG and microarray samples.

### Intracranial electrophysiology data

Awake intracranial electroencephalography data are obtained from the Open MNI iEEG atlas project^25^. This data includes 200Hz timeseries from 1772 channels from 106 patients (52 females, 54 males, Age=33.1 ± 10.8 years) with focal epilepsy. The timeseries are recorded from cortical grey matter and evaluated as normal by expert electrophysiologists according to strict clinical and research criteria. We retain channels from the left hemisphere of patients, compute the relative power of the alpha frequency band (8-12Hz) from the median power spectral density graph derived from the spectrogram of corresponding timeseries. Channels with z-scored values above 3 or below −3 are excluded. We co-register the remaining 836 channels to the MNI ICBM152 nonlinear 6th generation asymmetric template and interpolate relative bandpower to the reduced midthickness mesh.

### Serotonergic gene expression data

Gene expression levels for serotonin receptor HTR1A are obtained from the Allen Human Brain Atlas’ microarray expression data collected from six donors (1 Hispanic female, 2 African American males, and 3 Caucasian males; Age=42.5 *±* 12.2 years) and processed to retain probes with high differential stability (*>* 0.1) across donors using the abagen interface^35,53^. Retained probes are mapped to the MNI ICBM152 nonlinear symmetric 2009c template^100^, then transformed into fsLR space (equivalent to MNI ICBM152 nonlinear 6th generation asymmetric template) for interpolation to the reduced midthickness mesh. The nonlinear volume transforms are computed using ANTs.

### Surface and volumetric regression covariates

Regression covariates are used for testing the Spatially-Weighed Regression and Regression Kriging interpolation methods which include a linear regression step. For GRF data, we engineer toy covariates that are highly correlated with the ground truth by (i) adding noise from a uniform distribution with bounds −2 and 2, (ii) filtering along a single spatially-agnostic dimension, applying 3000 random spherical rotations and selecting the highest-correlated to the ground truth, weighing the GRF instance by a spherical Gaussian kernel with unit-variance centered at a random point, and (v) thresholding values below zero (Figure S1a). For empirical surface data, we select regression covariates by correlating all 24 maps and choosing five maps with the largest magnitude of correlation to the ground truth dense map (Figure S1b). For iEEG and microarray data that exist in volume as opposed to on the surface, we select regression covariates by exploring a set of 32 publicly available volumetric maps registered to the MNI ICBM152 nonlinear 6th generation asymmetric template (Figure S1c).

### Distance-dependent cross-validation

To establish reliability of variogram modeling and spatial interpolation for iEEG and microarray gene expression, we create a cross-validation sample using a distance-dependent approach (Fig. S6). We begin by seeding a Gaussian kernel at the geodesic centroid of each parcel of the Desikan-Killiany atlas and take five same-sized subsamples of either the sparse iEEG or microarray data, without replacement. This procedure is repeated for all 35 regions of the Desikan-Killiany atlas including the medial wall, yielding 175 samples each accessing 20% of the original sparse sample with a different spatial coverage and density. This sampling strategy ensures that for every seed region from the atlas, training data is non-overlapping.

## Acknowledgements

We thank A. Farahani, E. G. Ceballos, F. Milisav, M. Pourmajidian, T. Taheri, and M. Haddad for their comments and suggestions on the manuscript. We also thank A., A., and A. Pearson for helpful discussions. Y.Z. acknowledges support from the Natural Sciences and Engineering Research Council of Canada (NSERC), the Faculty of Medicine and Health Sciences of McGill University, the FSB Miller Fund, and the Kaplan Family Fund. B.M. acknowledges support from the NSERC, the Canadian Institutes of Health Research (CIHR), the Brain Canada Foundation Future Leaders Fund, the Canada Research Chair Program, the Michael J. Fox Foundation, and the Healthy Brains for Healthy Lives initiative. The funders had no role in study design, data collection and analysis, decision to publish or preparation of the manuscript.

---

**Figure S1.**
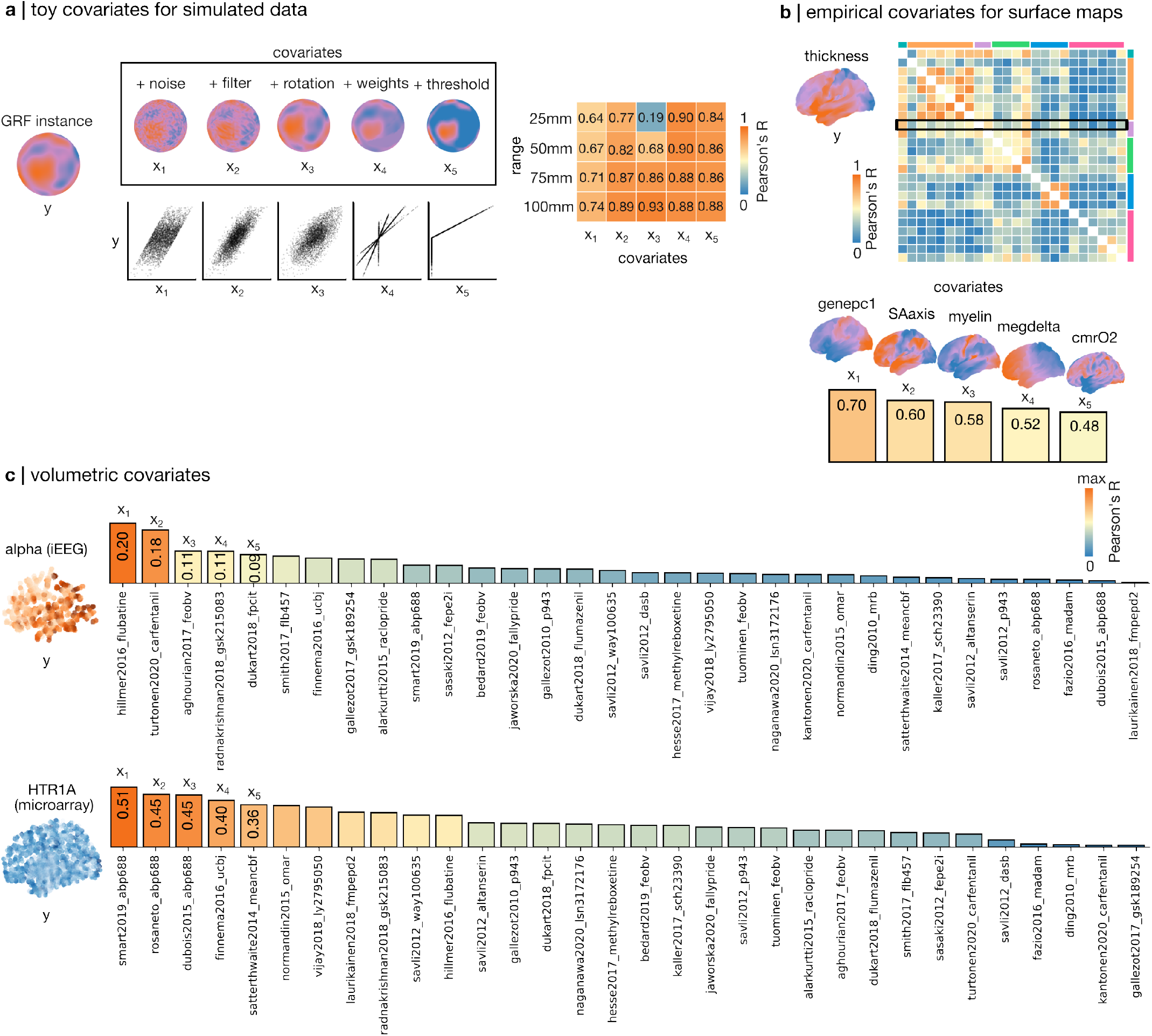
Regression covariates for spatial interpolation. Regression covariates *X*_*m*_ = [*x*_1_, *x*_2_, …*x*_*m*_] are necessary for interpolation with Spatially-Weighted Regression and Regression Kriging. For each scenario for *y* (GRF, surface maps, and volumetric sparse data) we include five covariates *x*_1_, *x*_2_, …, *x*_5_. **(a)** Toy covariates for GRF instances are created from the original data, by adding random noise, applying a Gaussian filter along a single spatially-agnostic dimension, applying a randomly determined (small) angular rotation, weighing the GRF with a Gaussian kernel, and thresholding for positive values only. Separate sets of toy covariates are made for GRF instances with range 25-100mm, and their correlation with *y* is overall high (*R >* 0.50). **(b)** Regression covariates for brain surface maps are empirically selected as the top five most correlated maps from a set of 24 empirical surface maps. The absolute correlation values are overall weaker for a number of empirical surface maps (*R <* 0.5). **(c)** Regression covariates for volumetric use cases are empirically selected as the top five most correlated volumetric images at sampled locations from a set of 32 empirical volumes. The absolute correlation values are very weak between imaging volumes and intracranial EEG (*R <* 0.2), and low-to-medium between microarray gene expression and the same imaging volumes (best *R* ≈ 0.5).

**Figure S2.**
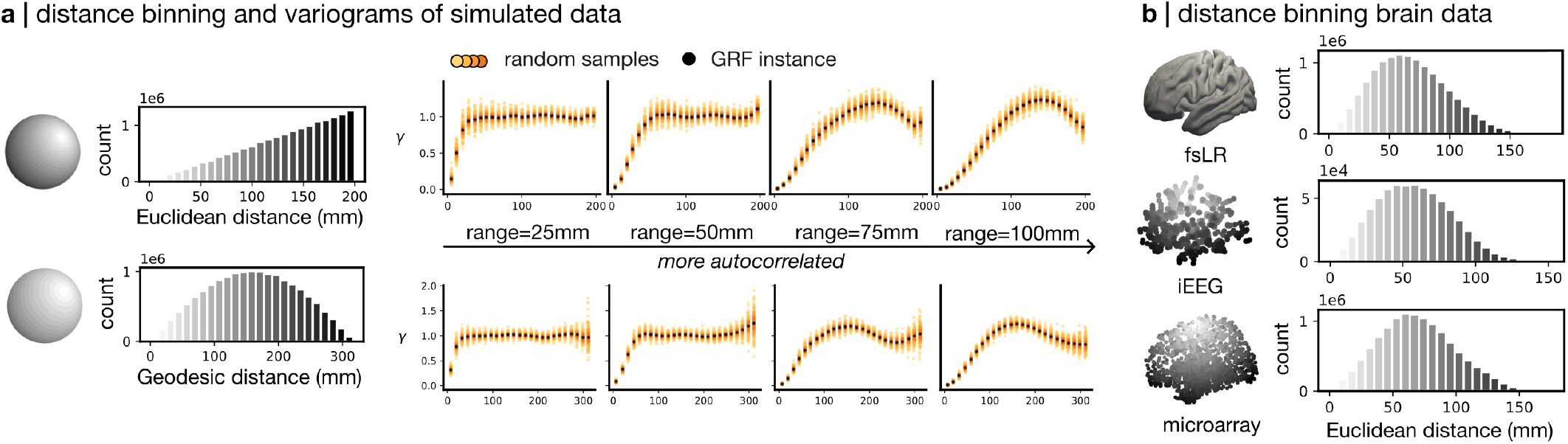
Distance-binning and variography. Experimental variograms are calculated by averaging semivariance (*γ*) values between pairs of points or vertices within distance bins. Across all geometries tested (sphere, fsLR brain surface, iEEG coverage, and microarray probe coverage), we conduct variography using 25 even distance bins. **(a)** On the spherical mesh, Euclidean distance bins oversample long-range pairs while geodesic distance bins oversample mid-range pairs. However, the shape of the experimental variogams adopt a similar transition from being approximately Gaussian to being cyclical. Geodesic distance-based estimates of semivariance is more unstable than Euclidean distance-based estimates at long ranges. **(b)** On a brain mesh or point set, Euclidean distance bins oversample mid-range pairs. Modes are within distance bins 65.6-75.0mm (fsLR), 49.0-55.0mm (iEEG), and 49.5-56.5mm (microarray). Euclidean distance is used for all the spatial models examined in this manuscript because it is compatible with all parametric variogram models.

**Figure S3.**
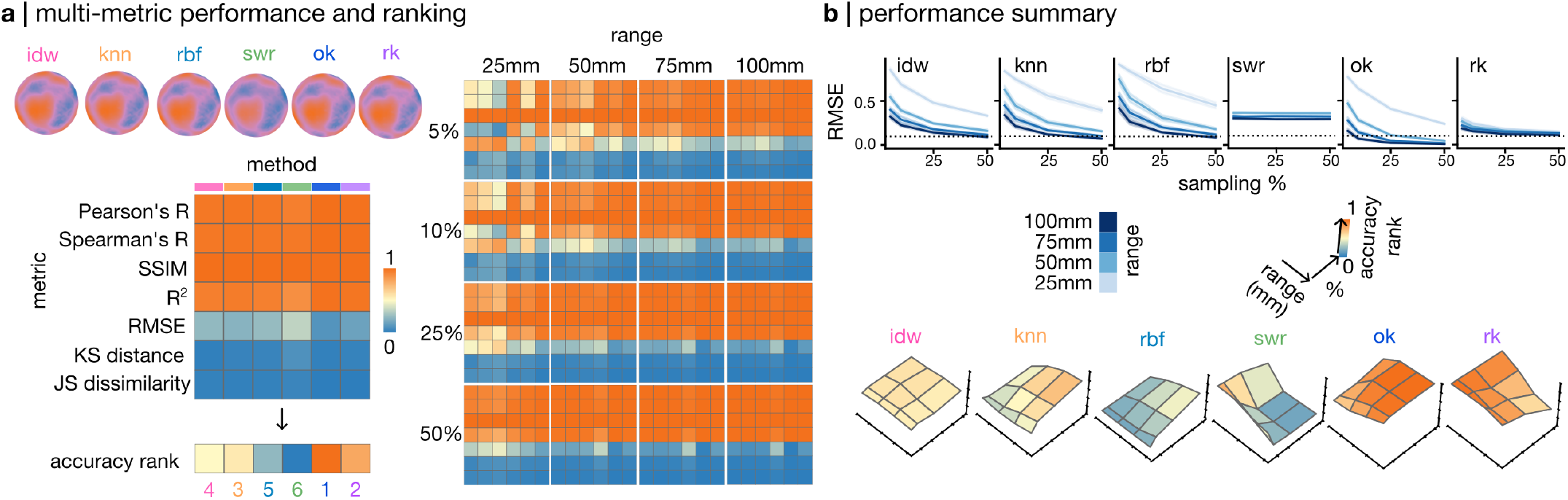
Ranking performance with spherical simulations. Dense maps interpolated from sparse subsamples are bench-marked against a ground truth dense map for the scenarios with GRF. We show examples of Inverse Distance Weighting (idw), K-Nearest Neighbours (knn), Radial Basis Function (rbf; Gaussian kernel), Spatially-Weighted Regression (swr), Ordinary Kriging (ok) and Regression Kriging (rk) interpolation for a GRF instance with range 50 sampled at 25%. **(a)** Seven similarity metrics are computed between an interpolated map and the ground truth at interpolated locations. These values capture the absolute performance of each method. Higher values for Pearson and Spearman correlation coefficient, structural similarity index, and *R*^2^ promote rank; higher values for root mean squares error (RMSE), Kolmogorov-Smirnov (KS) distance, and Jensen-Shannon (JS) dissimilarity demote rank. An aggregate rank score is computed from the average of individual rank scores. A higher rank score means that the interpolated map is, across all aspects of similarity, a more successful reconstruction of the ground truth. For GRF instances with ranges 25, 50, 75 and 100, and random subsamples at 5, 10, 25 and 50% of the mesh, we show absolute performance averaged across 100 runs for each range-percentage combination. Sparser samples with shorter range (smaller spatial autocorrelation) are more difficult to interpolate, in which case Spatially-Weighted Regression and Regression Kriging readily outperform the rest of the methods when provided with regression covariates that are strongly correlated to the ground truth (see Figure S1a). **(b)** Learning curves for each spatial interpolation method with confidence intervals established by 100 random samples for each range. The dotted line marks *RMSE* = 0.1 (99% variance explained). Accuracy rank scores are averaged across 100 runs for each range-percentage combination, summarizing accuracy across different benchmark metrics.

**Figure S4.**
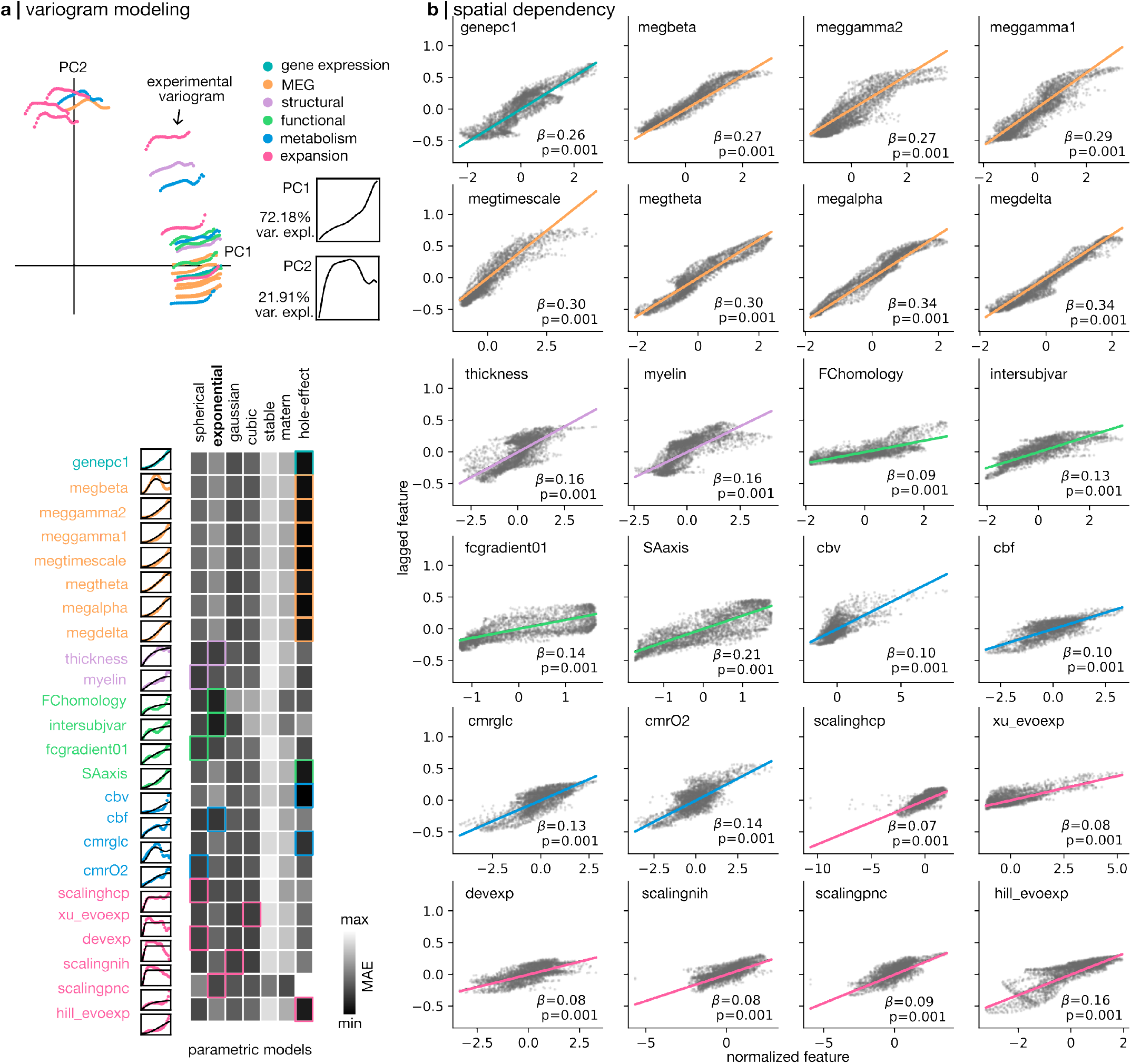
Spatial modeling for surface brain features. Each empirical surface maps contains a different spatial covariance structure. **(a)** We decompose the experimental variograms of each feature map using functional principal component analysis and retain the first two principal components (PC1, 2) that collectively explain 94.1% of variance. Different parametric variogram models are fitted to describe different features and we highlight the ones with the best fit. We include all positive-definite parametric models from Scikit-Gstat’s variogram repertoire, as well as a cyclical model (Bessel’s hole effect model) proposed by^S99^. **(b)** To assess the spatial dependency of empirical features maps, we visualize Moran scatterplots for every feature map^S5^. Briefly, the Moran scatterplot summarizes spatial dependency as the linear relationship between the feature values and lagged values (feature vector multiplied by the row-normalized distance weight matrix) of the underlying data. The slope *β* of this linear relationship is the Moran’s I statistic. We compute a p-value for the Moran’s I statistic using 1000 spatial permutations under the null hypothesis *β* = 0 (i.e., random spatial lags).

**Figure S5.**
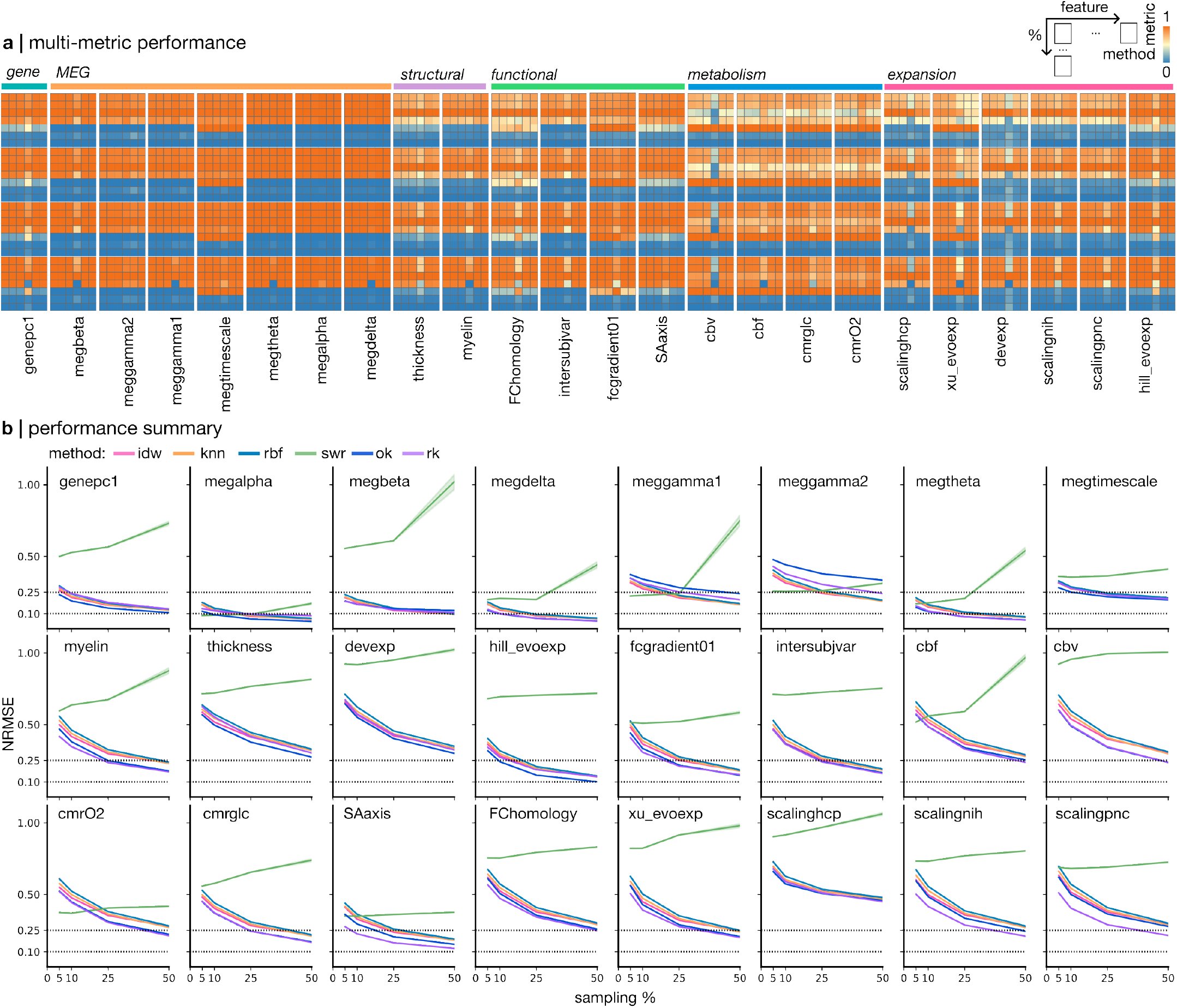
Ranking performance with empirical surface maps. **(a)** For all empirical surface brain maps, we show absolute performance averaged across 100 runs for each feature-percentage combination, sampled using an anatomy-first approach. The rows of heatmaps represent the same sampling percentages, while the rows in the heatmaps are each a similarity metric. Each column of heatmaps is a brain feature, while the columns in the heatmaps are each an interpolation method. These metrics are used to produce the ranking shown in Fig. 5. Metrics and methods are ordered as in Fig. S3a. **(b)** Learning curves for each feature map, across all spatial interpolation methods with confidence intervals established by 100 anatomy-first subsamples taken at different percentages. Dotted lines mark *NRMSE* = 0.25 and 0.10, for 94% and 99% variance explained, respectively.

**Figure S6.**
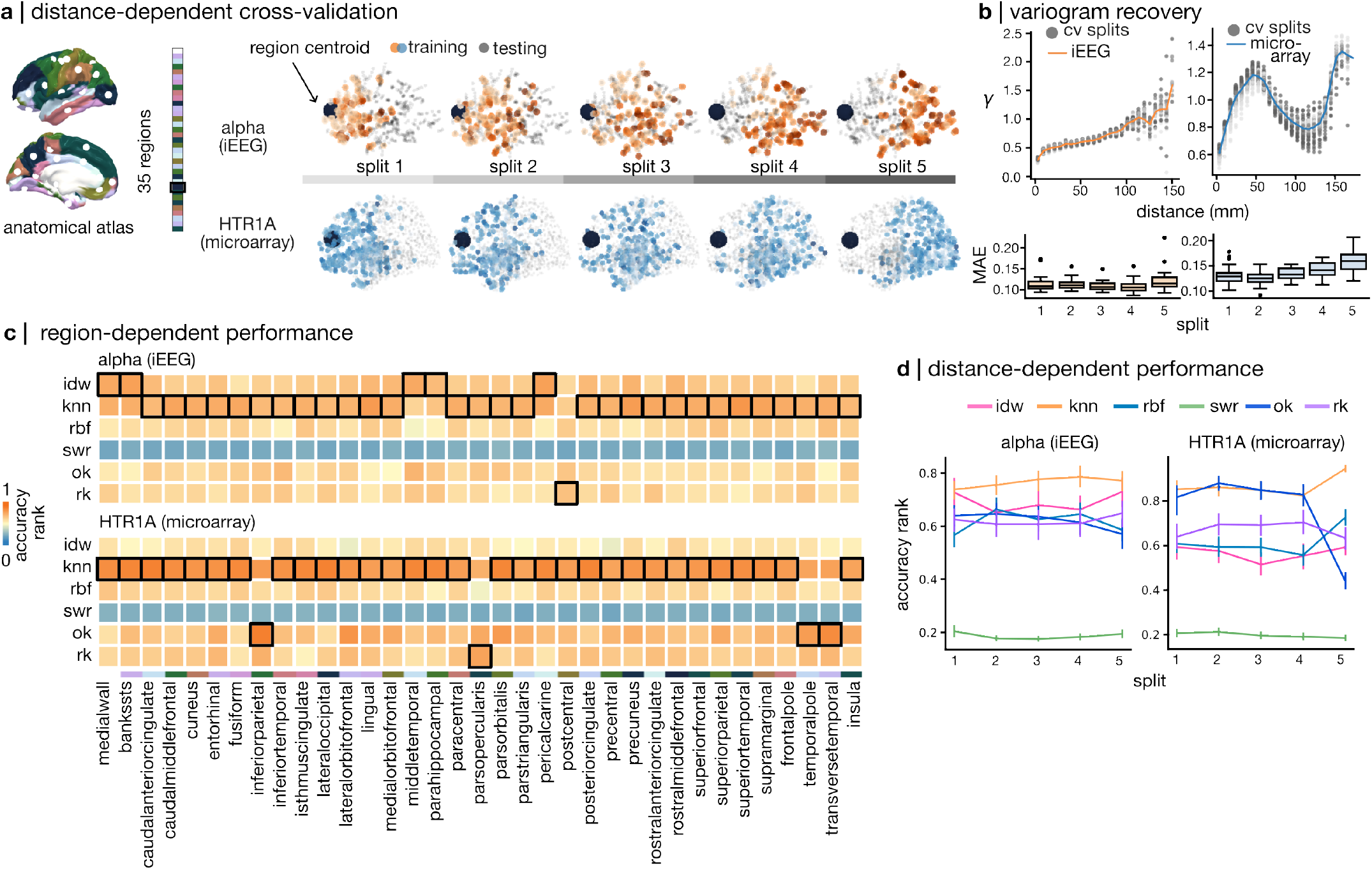
Distance- and region-dependent cross-validation. For use cases with sparsely sampled volumetric brain data, we benchmark interpolation methods by criteria of internal validity using distance-dependent cross-validation. **(a)** At the geodesic centroid of each region of the Desikan-Killiany anatomical atlas, we center a Gaussian kernel (sigma set to the 0.2-th quantile of distances from the centroid) and split the data into five non-overlapping subsets to interpolate from. Each split samples 20% of the original sparse data. **(b)** Experimental variograms of cross-validation (CV) subsamples as well as the original data are computed using the same binning scheme. The CV variograms are coloured by split (greyscale). We fit an exponential variogram model to experimental variograms derived from each CV subsample and obtain a mean absolute error (MAE) value for the fit. **(c)** Accuracy ranks are scored across atlas regions and averaged across all five splits per region. The best-scoring interpolation method per region is boxed. **(d)** Accuracy rank for each method varies across CV splits. Confidence intervals are established by 35 CV samples per distance-dependent split.

